# Task specificity in mouse parietal cortex

**DOI:** 10.1101/2020.12.18.423543

**Authors:** Julie J Lee, Michael Krumin, Kenneth D Harris, Matteo Carandini

## Abstract

Parietal cortex is implicated in a variety of behavioral processes, but it is unknown whether and how individual neurons participate in multiple tasks. We trained head-fixed mice to perform two visual decision tasks involving a steering wheel or a virtual T-maze, and recorded from the same parietal neurons during the two. Neurons that were active during the T-maze task were typically inactive during the steering-wheel task, and vice versa. Recording from the same neurons in the same apparatus without task stimuli yielded the same specificity as in the task, suggesting that task specificity depends on physical context. To confirm this, we trained some mice in a third task combining the steering wheel with the visual environment of the T-maze. This hybrid task engaged the same neurons as the steering-wheel task. Thus, participation by neurons in mouse parietal cortex is task-specific, and this specificity is determined by physical context.

## Introduction

The brain must meet a vast variety of potential behavioral demands while relying on a finite number of neurons. This behavioral diversity would be challenging if all neurons had a single, fixed function. Although classical studies of the early sensory cortex suggested individual neurons map to a specific sensory feature arising from a single sense organ, studies of higher cortex, and more recent investigations of primary visual cortex reveal a more complex picture (e.g. Stringer et al., 2019; Musall et al., 2019). Further, neurons in many regions exhibit “mixed selectivity”, firing for multiple aspects of behavior (Rigotti et al 2013; Parthasarathy et al 2017; Mante et al 2013; Raposo et al 2014, Park et al. 2014; Meister et al. 2013; Zhang et al 2017). Mixed selectivity typically has been studied during a single behavioral task, but suggests that neurons might likewise participate in multiple tasks.

A region where one might expect to find neurons involved in multiple behavioral tasks is the parietal cortex. Parietal neurons have been implicated in many aspects of vision, decision, and navigation, including motor planning (e.g. Gnadt and Andersen 1988; Snyder et al. 1997), evidence accumulation (e.g. Britten et al. 1996; Shadlen and Newsome 2001; Hanks et al. 2015; Pinto et al., 2019) choice sequences (Harvey et al. 2012), spatial position and heading (Krumin et al., 2018), movement motifs (Chen et al. 1994; Wilber et al. 2014; Whitlock et al. 2012), and movement sequences (Nitz, 2006; Nitz et al. 2012). Parietal neurons can exhibit mixed selectivity for combinations of these behavioral variables within a single task (Parthasarathy et al 2017; Mante et al 2013; Raposo et al 2014; Park et al. 2014; Meister et al. 2013; Zhang et al 2017).

It is not clear however, how individual parietal neurons participate in multiple tasks. Experiments testing this question are rarer, because they require training subjects on multiple tasks and recording from the same individual neurons across them. A study in primates found that parietal neurons participate in two versions of a visual task based on color or motion (Mante et al 2013). Likewise, studies in rodents found that parietal neurons have similar responses when a given behavioral report was based on visual vs. auditory stimuli (Raposo et al., 2015) or visual vs. tactile stimuli (Nikbakht et al., 2018). On the other hand, a study in primates that employed different means of behavioral reports found a strong segregation of parietal neurons, with neurons in one area responding when the report involved eye movements and neurons in a nearby area responding when it involved arm movements (Snyder et al., 1997). It is not clear whether and how these results can be reconciled; perhaps certain aspects of these task designs determine participation in a task.

Here we recorded from large populations of parietal neurons while mice performed two visual decision tasks in different sensorimotor contexts. Both tasks required a two-alternative forced choice to indicate the presence of a grating stimulus, but the tasks used different visual stimuli, required different motor outputs, and used different physical experimental apparatuses. In the first task, mice walked on an air-floating ball to navigate in a virtual T-maze (Krumin et al., 2018). In the second task, mice turned a steering wheel in a visual contrast-detection experiment (Burgess et al., 2017; International Brain Laboratory, 2020). Surprisingly, the two tasks activated largely distinct but spatially-intermixed subpopulations of neurons, and the few neurons that were activated by both did not show similarities in their responses across the two tasks. Additional experiments established that this specificity was driven by the physical context of the experimental apparatus. Individual neurons in parietal cortex thus are not generalists but are rather specialists, active only in specific contexts.

## Results

We trained mice to perform two visual decision tasks while head-fixed (Fig. 1a,b). In the first task (“T-maze task”), mice ran on an air-suspended styrofoam ball to navigate through a virtual T-maze, and reported the location of a grating present on the left or right wall by turning into the corresponding arm at the end of the initial corridor (Fig. 1a, top) (Krumin et al., 2018). In the second task (“steering-wheel task”), mice sat on a platform and turned a steering wheel with their front paws, reporting the location of a grating on the left or right side by turning the wheel to bring the grating to the center (Fig. 1b, top) (Burgess et al., 2017). In both tasks, the visual contrast on each trial was chosen from a range of values to vary difficulty. We trained six mice to perform both tasks consecutively in the same day. Mice typically performed hundreds of trials in each task. They exhibited good performance in both tasks, making more rightward choices with higher contrasts for stimuli on the right and more leftward choices with higher contrasts for stimuli on the left (Fig. 1a,b, bottom). They seldom made mistakes when stimuli had high contrast, and chose randomly (nearly 50-50) between left and right when stimuli were absent, i.e. at zero contrast.

**Figure 1.**
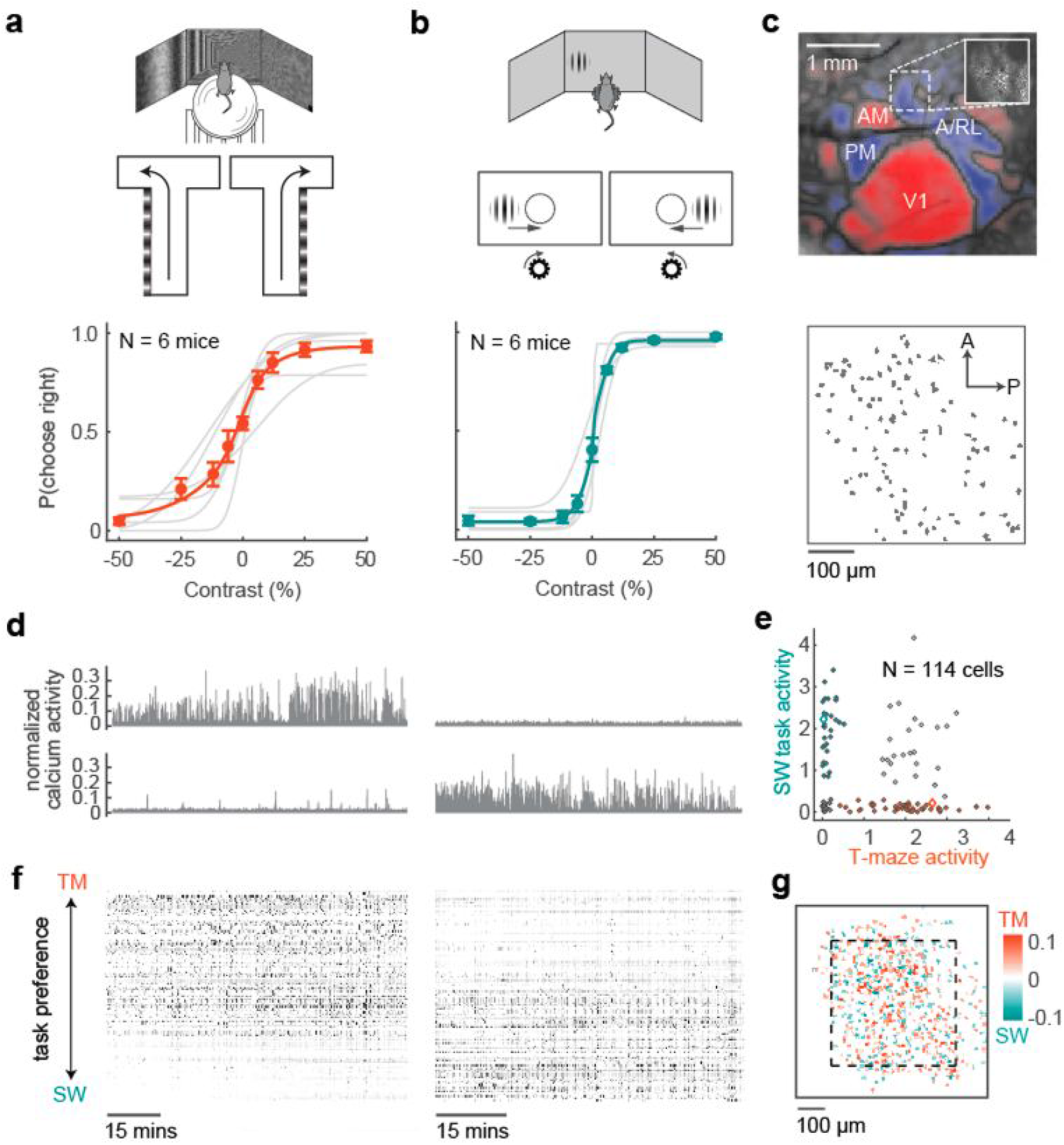
In mice performing two visual decision tasks, many parietal neurons are task-specific. (a) The T-maze task. Mice walk on an air-suspended ball to navigate a virtual corridor with a grating painted on one wall, turning left or right at the end of the corridor to indicate the grating position (*top*). *Bottom*: performance of 6 mice in the task, plotting fraction of rightward choices vs. contrast of stimuli on the left (negative) or on the right (positive). Dots and error bars show mean ± s.d. for N = 21 sessions in 6 mice. Curves show the fitted psychometric function for each mouse (*gray*) and averaged across mice (*orange*). (b) The steering-wheel task. Mice turn a steering wheel to indicate whether a grating is on the left or right (*top*). *Bottom*: performance in the task of the same mice on the same days as (a). Psychometric curves for individual sessions are in Supplementary Figure S1. (c) *Top*: map of visual cortical areas from widefield imaging, showing the visual field sign of retinotopic areas (*blue:* negative; *red*: positive) and the field of view targeted for two-photon imaging (*inset*) from an example mouse. Bottom: outlines of the identified neurons in the field of view. (d) Responses of two neurons from the example session, showing task-specific activity. (e) Summary of activity (isolation distance) of the 114 neurons in the example session in the T-maze (TM) vs. steering-wheel (SW) tasks, showing neurons that fired only in the T-maze task (*orange*), only in the steering-wheel task (*blue*), in both tasks (*white*), or in neither task (*gray*). *Diamonds* indicate the example neurons in (d). (f) Raster plot showing the firing of neurons in an example session in the two tasks. Gray level denotes deconvolved calcium signal, z-scored. Neurons are sorted by relative task preference for the two tasks (the diagonal of (e)). (g) Anatomical distribution for example mouse from (c-f) showing the overlay of ROIs over 9 sessions, colored as in (e). Dashed square indicates a typical imaging field of view as in (c).

We then used two-photon calcium imaging to record from the same population of parietal neurons in the two tasks (Fig. 1c). We targeted a parietal region anterior to the primary visual cortex and overlapping with visual areas A and RL (Hovde et al., 2018, Wang et al., 2020; Gilissen et al., 2020), identified by mapping retinotopy with widefield imaging (Zhuang et al, 2017; Garrett et al., 2014; Sereno et al., 1994; Fig. 1c, top). We targeted this region because it is readily identified and distinguished from nearby somatosensory and primary visual areas. We then imaged this region with a two-photon microscope (Fig. 1c, inset) to record the activity of hundreds of parietal neurons simultaneously (Fig. 1c, bottom). Mice were tested on both tasks in the same imaging rig.

Parietal neurons could participate in either task, but were typically task-specific (Fig. 1d-g). Neurons that were active during the T-maze task were typically inactive during the steering-wheel task, and vice versa (Fig. 1d). To quantify this effect, we summarized the activity of each neuron within each task using the “isolation distance” measure (Stringer & Pachitariu, 2019), which characterizes a neuron’s activity level relative to background neuropil fluorescence. Comparing this measure across tasks revealed a large fraction of neurons that were active only in the T-maze or only in the steering-wheel task (Fig 1e, orange and blue), but only a few active in both tasks (Fig 1e, gray). Similar results were obtained using other measures of activity (Supplementary Figure S2). We could then use the difference in activity across tasks to sort the neurons in order of task preference, illustrating consistent differences in participation across tasks: most neurons fired during one task but rarely in the other (Fig. 1f). Task-specific neurons seemed to intermix, with no obvious anatomical organization (Fig. 1g).

Recording the same population across days revealed that this task specificity was robust and repeatable (Fig. 2). To record from the same neurons across days, we imaged the same plane on a subsequent day and aligned cells recorded on both days using Suite2p (Pachitariu et al., 2018). We then compared each neuron’s activity across days, within or across tasks (Fig. 2a). Activity across days was highly correlated within tasks (Fig. 2b) but negatively correlated or not significantly correlated across tasks (Fig. 2c). Activity was significantly more similar within than across tasks (Fig. 2d), whether considered for the T-maze (p = 9e-6) or steering-wheel task (p = 2e-4, one-tailed t-tests). These results indicate that the task specificity shown by many parietal neurons is robust and largely stable across successive days (Fig. 2e).

**Figure 2.**
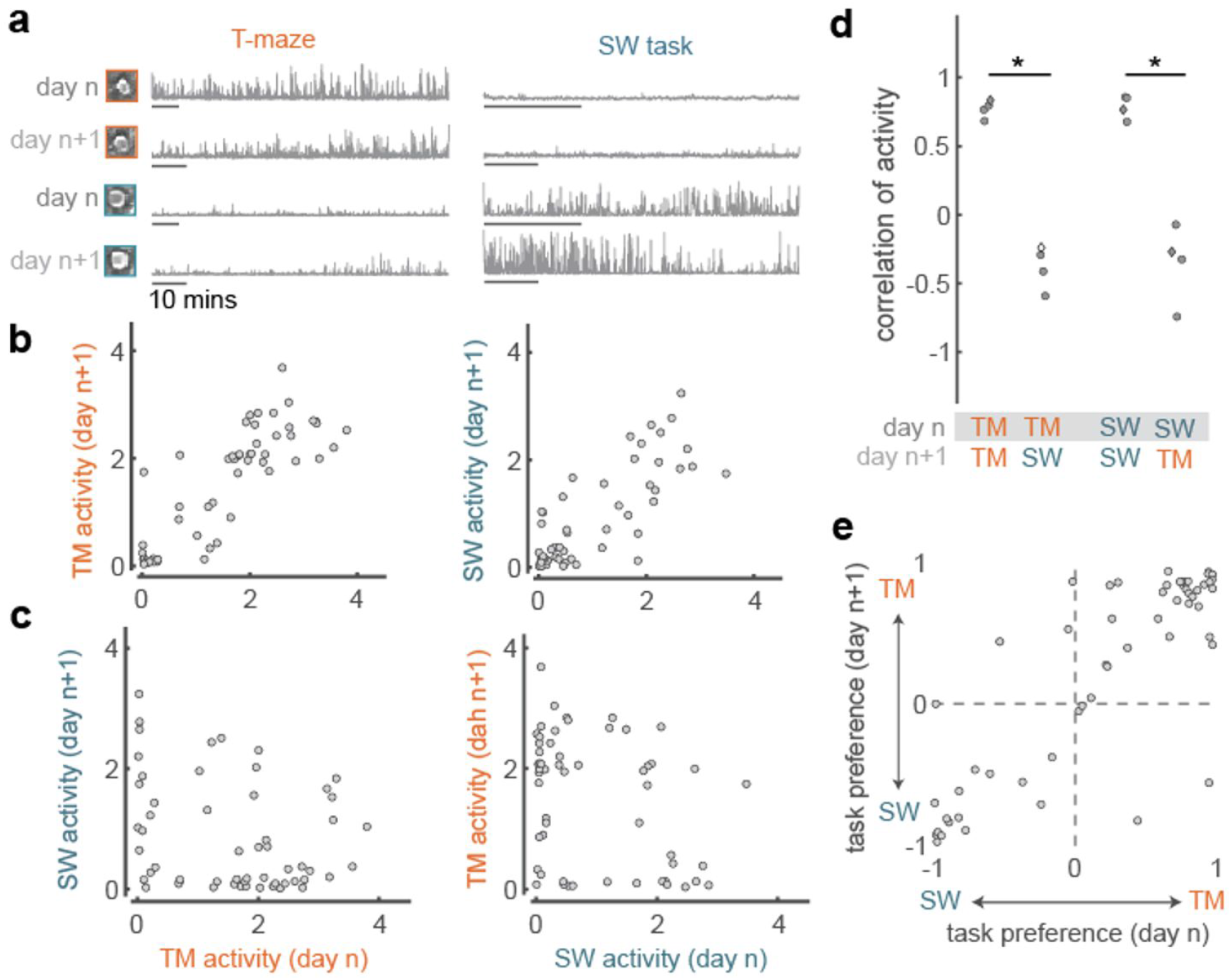
Task-specificity is consistent across days. (a) Activity of two example neurons in the T-maze on consecutive days (*left*). Activity of the same neurons in the steering-wheel task across days *(right*). Insets show the mean images of each neuron in each session. Each bar represents 10 minutes. (b) Comparison of activity within tasks across consecutive days, in the T-maze (*left*) or steering-wheel task (*right*). Correlations were positive in both cases (r = 0.83 and r = 0.77, p ≈ 0, i.e. too small to measure). (c) Same as in (b) but comparing activity across tasks. Correlations were negative (*left*: r = −0.24, p = 0.08) or not significant (*right*: r = −0.27, p = 0.05). (d) Summary from four pairs of days in three mice. *Diamond* illustrates the example pair of days from (b-c). Filled points indicate significant Spearman rank correlations at p < 0.05. (e) Comparison of task preference (relative activity over tasks: positive for neurons preferring the T-maze task and negative for neurons preferring the steering wheel task) for neurons imaged in two example consecutive days (N = 56 cells), showing significant correlation across days, r = 0.84, p = 5e-16. Correlations were also high in the other three pairs of days, with r = 0.85, 0.87, and 0.78.

The task specificity of parietal neurons must then be attributable to repeatable factors that are inherent to each task. The distinguishing factors might lie in the sensory context: though both tasks are based on vision, one involves visual scenes in virtual reality and the other involves a spatially-isolated visual grating. The distinguishing factors may also lie in the physical context: the apparatus used to perform each task (an air-suspended ball vs. a steering wheel), or the associated motor demands (running vs. steering).

To investigate the role of physical context, we recorded the same neurons in each task apparatus while mice passively viewed a gray screen, and found that neurons had similar specificity as in the task (Fig. 3). In these passive conditions, the firing of neurons was similar to the firing in the task corresponding to the same apparatus, and different from the firing in the other apparatus (Fig. 3a). Across these two behavioral conditions (task vs. passive) activity was highly correlated within a physical context (Fig. 3b), and uncorrelated or negatively correlated across contexts (Fig. 3c). Correlations were significantly different across contexts but not within contexts (Fig. 3d, one-way ANOVA, F(3,36) = 9.43, p = 1e-16). This role of context was not explained by the presence or absence of running, which is possible only on the ball: neurons’ task-specificity did not correlate with their modulation by running speed (median r = 0.09 +/- 0.09; p > 0.05 for 10/10 sessions, Supplementary Figure S3).

**Figure 3.**
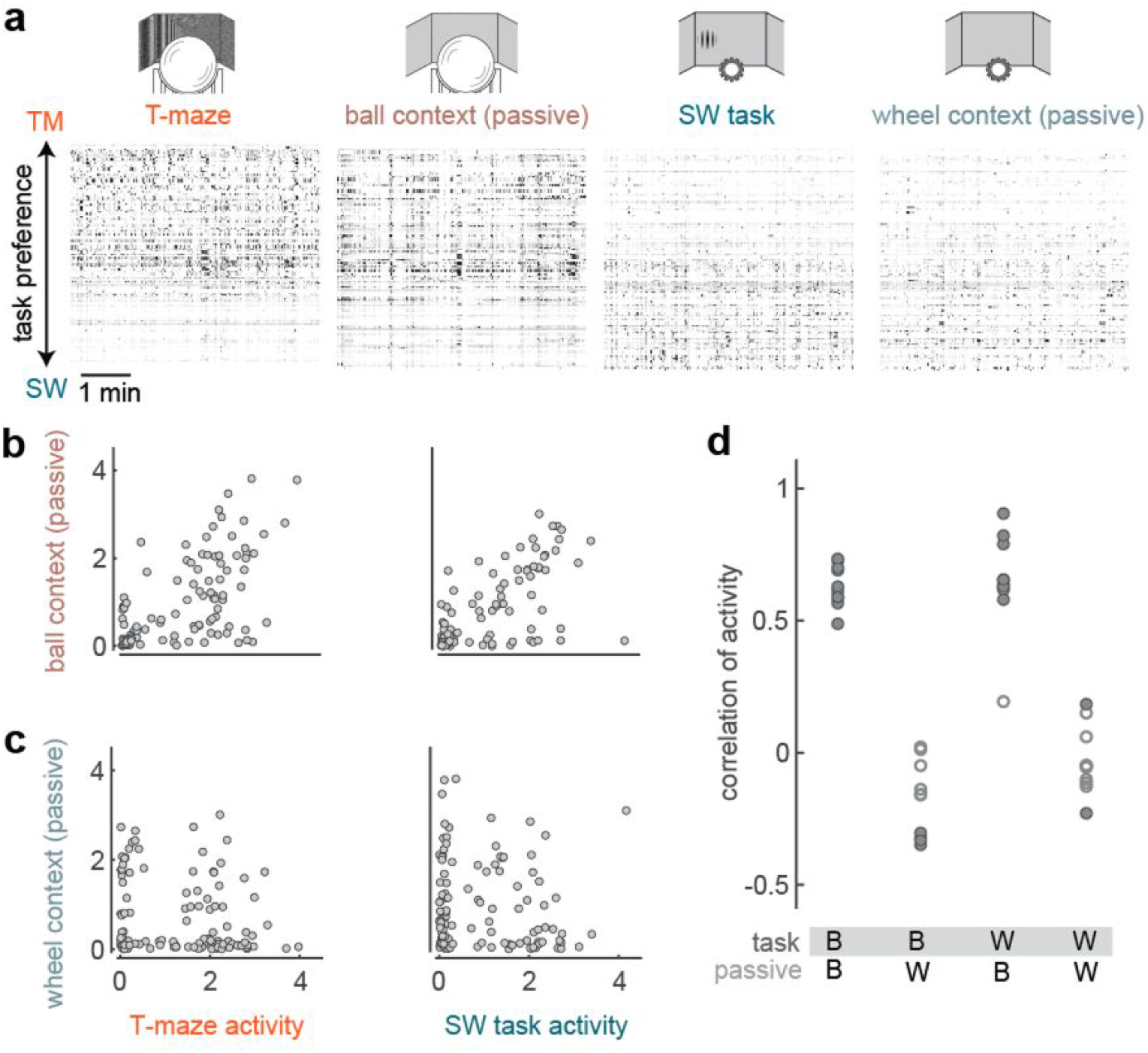
Task-specificity is predictable by physical context in the absence of a task. (a) Raster plot of activity from neurons in an example session showing five-minute segments of activity in each task and in the corresponding passive condition. Gray level indicates normalized firing rate as in Fig 1. Left to right: T-maze, passive ball, steering-wheel task, passive steering-wheel. (b) Comparison of activity for the same population of neurons across conditions with similar physical context, for the example session in Fig 2a-c. Activity is highly correlated both within the ball context (*left*, r = 0.63, p ≈ 0) and within the wheel context (*right*, r = 0.65, p ≈ 0) (c) Comparison of activity across different physical contexts for the same session. Activity is not significantly correlated (*left*: r = −0.16, p = 0.09; *right* r = −0.10, p = 0.28). (d) Summary of correlations of activity within and across physical contexts for 10 sessions where we recorded passive conditions (B = ball context; W = wheel context). Filled circles indicate significant Spearman rank correlations.

To further confirm the role of physical context in determining task specificity, we trained two of the mice in a third, hybrid task, which combined the visual context of the T-maze with the physical context and motor demands of the steering wheel (“steering T-maze” task, Fig 4a). Mice viewed the virtual T-maze environment from a fixed position in the middle of the corridor and turned the steering wheel to change the view angle (Fig. 4a). To report a choice, mice were required to turn the steering wheel so as to orient towards the wall with the grating. The gain of the steering wheel matched the steering-wheel task, such that a choice required the same amount of turning in both tasks.

**Figure 4.**
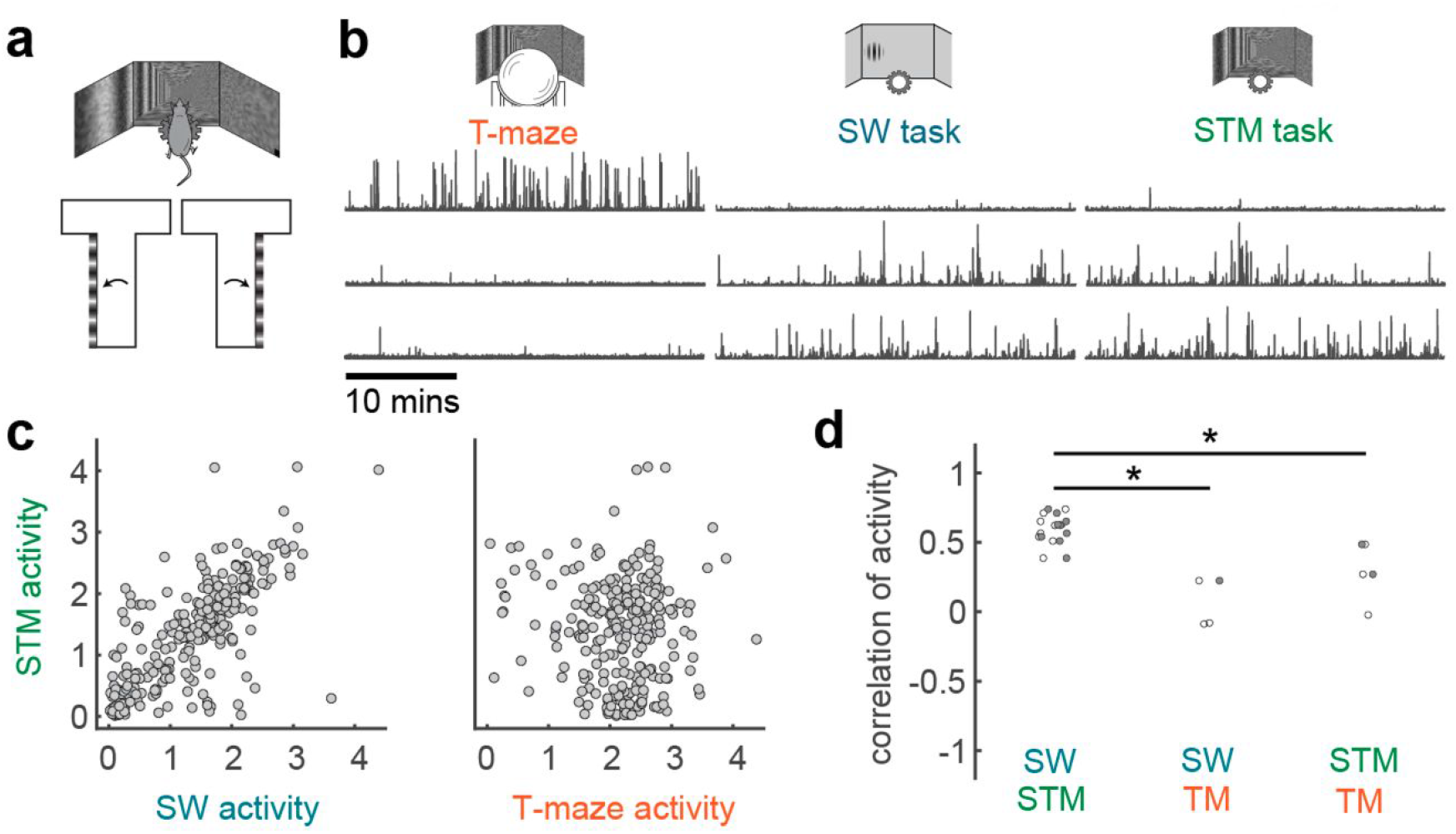
Activity in a hybrid task confirms the role of physical context. (a) The “steering T-maze” task (STM) combines the apparatus of the steering wheel with the visual scene of the T-maze in a fixed position along the corridor. (b) Three example neurons from a session that included all three tasks. (c) Activity of the same population of neurons across the steering-wheel and steering T-maze tasks (*left*, r = 0.74, p ≈ 0) and across the T-maze and steering T-maze tasks (*right*, r = −0.02, p = 0.72) from the same session as (b). (d) Summary of pairwise comparisons between the T-maze, steering-wheel, and hybrid tasks. SW vs hybrid: n = 9 sessions; SW vs TM: n = 3 sessions; TM vs hybrid: n = 3 sessions. We only included sessions in mice tested on the third task, and only compared pairs of tasks for which a sufficient number of trials was acquired. We performed an independent group means test (one-way ANOVA) as sample sizes were unequal, F(2,12) = 17.74, p = 0.0003.

The hybrid task engaged the same parietal neurons as the steering-wheel task, confirming that the specificity of parietal neurons across tasks is determined by physical context (Fig. 4b-d). In some sessions the mice were able to perform all three tasks consecutively, albeit with a smaller number of trials per task, as expected because of satiation. In one such session we were able to follow the same neurons across the three tasks, and found that activity was similar across tasks with the same physical context but dissimilar across tasks with different physical contexts (Fig. 4b-c). Similar results were seen in sessions where mice performed different pairs of the three tasks: as observed in the passive conditions, task participation was correlated within but not across contexts (Fig. 4d). These differences were highly significant (one-way ANOVA with unequal sample sizes, F(2,12) = 17.74, p = 0.0003). Therefore, the task specificity of parietal neurons is determined by physical context and not by visual context.

Having established that most parietal neurons are engaged only in specific physical contexts, we turned to the remaining neurons and asked if they carried similar signals across the two contexts. Parietal cortex has long been associated with decision making, and both of our tasks require the mouse to decide whether a grating is on the left or right. Neurons that participate in both tasks might thus encode choice signals shared across tasks. Since our tasks involve multiple visual contrasts, we calculated choice preference using an extension of “Choice Probability” (Britten et al., 1992) called “combined conditions Choice Probability” (ccCP; Steinmetz et al., 2019). This statistic measures the probability that a neuron’s firing rate was greater on trials with one choice than the other, for matched stimulus conditions (Methods).

Even in the neurons that were active across tasks, task-evoked responses bore little similarities across contexts (Fig. 5). Choice could be decoded from some neurons in each task (Fig. 5a,b), with choice encoding appearing to be common in the T-maze task (Krumin et al., 2018) and rare in the steering-wheel task (Steinmetz et al., 2019) (Fig. 5c). Choice preferences were correlated across days within the T-maze (Fig. 5a,b) but were not correlated across tasks (Fig. 5d-f). Therefore, even though these neurons had an opportunity to show similarities in the two tasks, because they were active in both, they likewise showed specificity in their responses.

**Figure 5.**
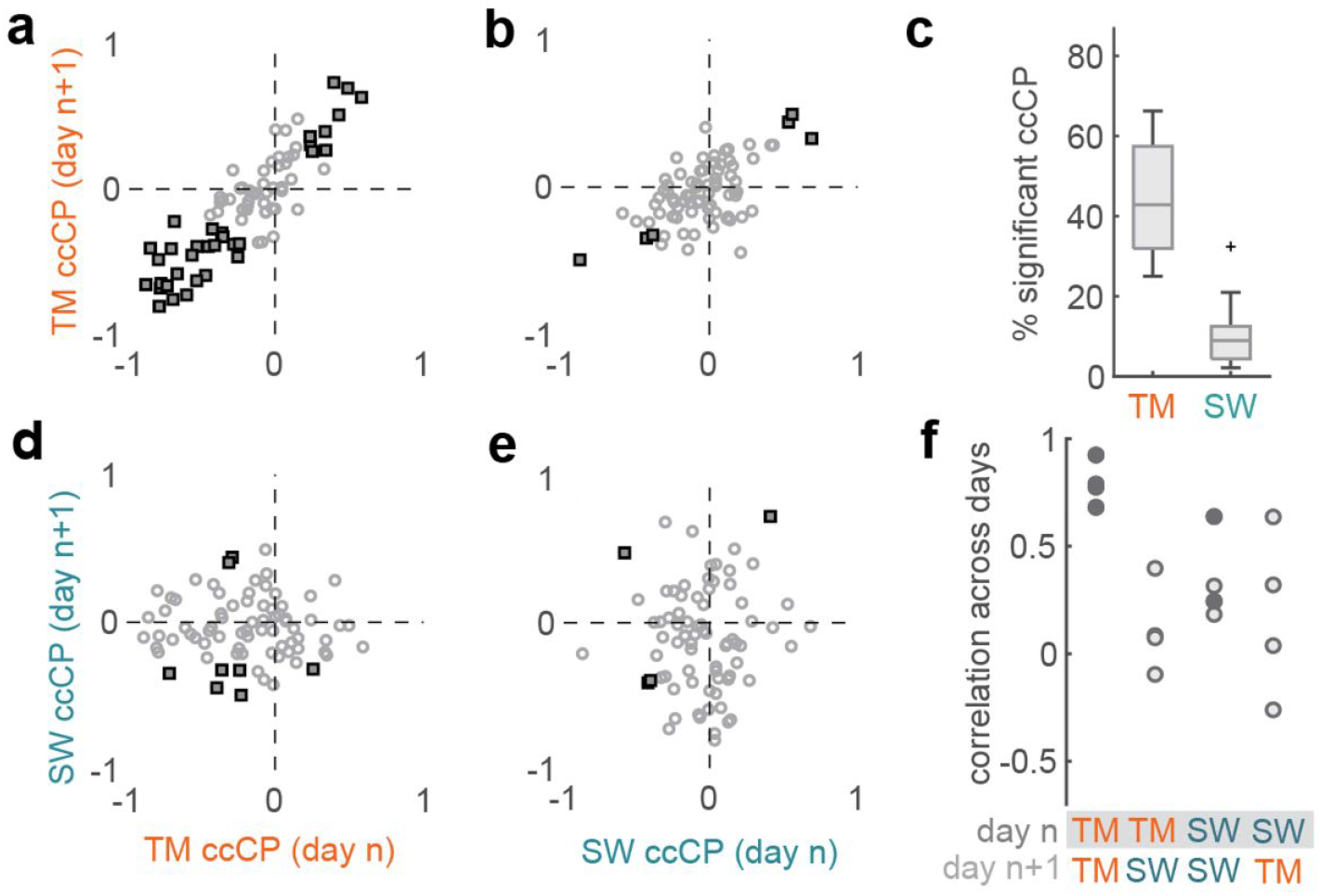
Preference for ongoing choice is not shared across tasks. (a) Choice-selectivity of neurons in the T-maze task, estimated across consecutive days using “combined conditions choice probability” (ccCP). Negative values denote preferences for left choices, positive for right choices. *Filled squares* denote neurons that were significantly selective for choice in either day (p<0.05). (b) Choice selectivity of the same population of neurons in the steering wheel task. (c) Proportion of neurons found to be significantly choice-selective over 21 sessions from 6 mice, for each task. Outliers are denoted by a *plus* symbol. (d,e) Comparison of choice selectivity across tasks. *Filled squares* indicate neurons that were significantly choice selective on both tasks. (f) Correlation of ccCP across days averaged across all neurons active in both tasks (n = 3 mice, four pairs of days, isolation distance > 0.3 in both tasks). *Filled circles* indicate a significant Spearman rank correlation.

## Discussion

By training the same mice in multiple visual decision tasks, we were able to assess how parietal neurons change their activity with behavioral demands. Most neurons were active during one task and inactive during the other. This task-specificity was reliable across successive days, indicating that it can be explained by factors inherent to each task. Indeed, recording in passive conditions and in a hybrid task established that the primary factor driving task-specific participation is the physical context. Physical context also influenced choice representations in the few parietal neurons that responded in both contexts: their choice preferences were not correlated across contexts. Therefore, we conclude that physical context is the dominant factor that determines whether parietal neurons participate in a given task and influences the content of their representations.

Our findings are consistent with the few studies that compared the activity of individual parietal neurons across tasks. In primate parietal cortex, different neurons were engaged for saccades and reaches (Snyder et al., 1997). Our results extend these observations: we were able to record from a large population of neurons across days, and established that task specificity is driven by physical context even in the absence of a task. Furthermore, this role of physical context is compatible with previous reports from rodent parietal cortex. In studies where rats switched between decisions based on different sensory modalities (Raposo et al., 2014; Nikbakht et al., 2018), subjects used the same motor action and behavioral apparatus to report choices, and neurons had correlated choice preferences across tasks. In our design, subjects used different motor actions to report choices, and most neurons did not participate in both tasks, while the few that participated in both tasks did not have correlated choice preferences across tasks.

Together with our results, these findings suggest that parietal neurons are task-general when tasks share the same physical context, and task-specific when the tasks differ in physical context. It may be that, in contexts where parietal neurons are active, they carry causal signals related to sensation or decisions (Zatka-Haas et al. 2020; Goard et al. 2016; Licata et al. 2017; Odoemene et al., 2018; Itokazu et al., 2017), but their participation is gated by physical context in a neuron-specific fashion. Also, it has been suggested that there is a tight relationship between choice and pre-motor encoding (Shadlen et al., 2008). Our results are consistent with this idea: because different motor actions are required in the two contexts, it is reasonable that choice signals would be conveyed by different neurons.

Overall, our findings emphasize the value of sampling multiple behaviors in the same neuronal populations. During any one task, most neurons are silent, but may be recruited when mice perform a different task. Presumably yet another population would have been active in a third physical context, raising the possibility that at least in parietal cortex, “silent cells” (e.g. Thompson & Best, 1989) might become active in an appropriate context. The activity of task-specific neurons when mice are placed in the appropriate apparatus, even without performing the task, also implies that “spontaneous” activity, at least in parietal cortex, is context-dependent: spontaneous behavior on an air-suspended ball engages one set of neurons, and spontaneous behavior on a steering wheel engages another set. Such context-specific spontaneous activity may reflect the mice’s knowledge of their current physical environment, as well as sensorimotor variables that differ between contexts, such as tactile cues or body posture. In general, the stark difference in activity that we observed between contexts might apply brainwide rather than restricted to parietal cortex, and might help place the brain in the appropriate state to perform the correct task in the correct context.

## Acknowledgements

We thank Laura Funnell, Miles Wells, and Hamish Forrest for assistance with training, and Charu Reddy for laboratory support. This work was supported by the Wellcome Trust through a PhD studentship to JJL (grant 109004) and Investigator Awards to KDH and MC (grants 205093 and 108726). MC holds the GlaxoSmithKline / Fight for Sight Chair in Visual Neuroscience.

## Author Contributions

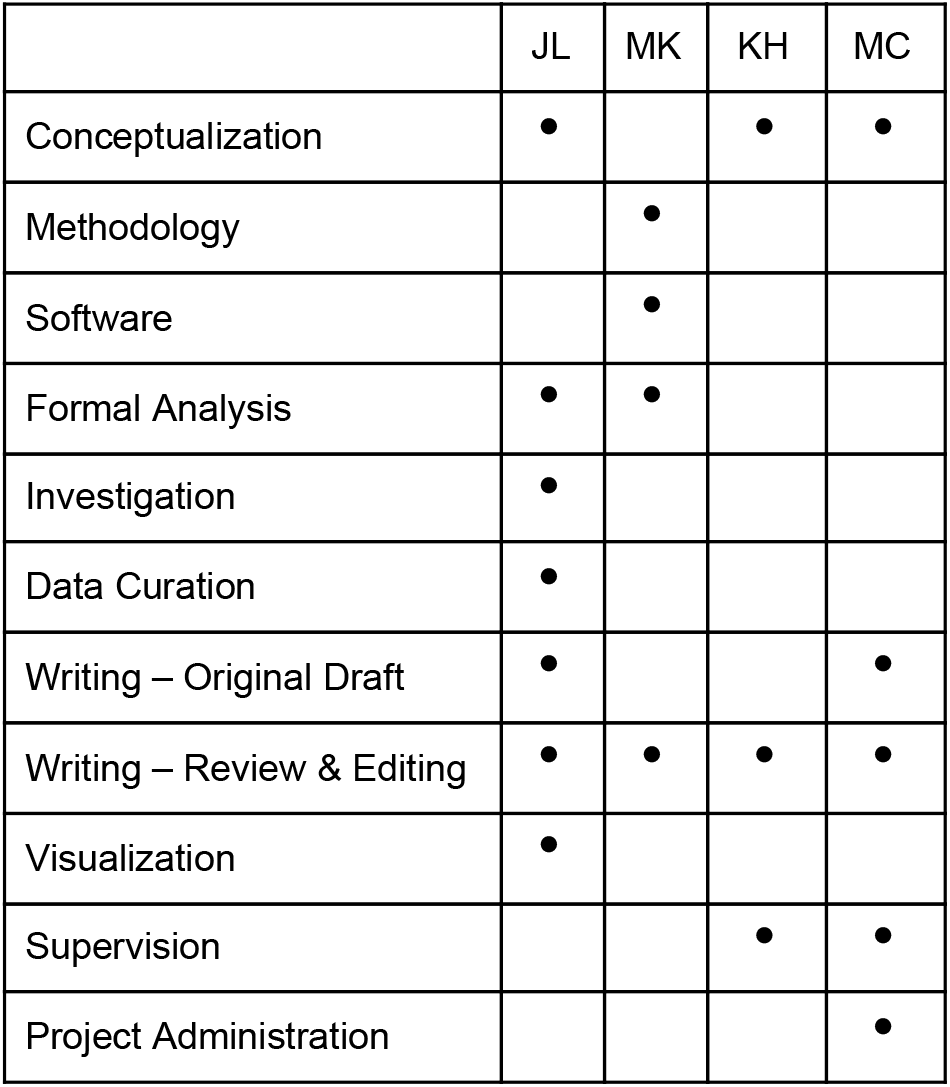

## Declaration of Interests

The authors declare no competing interests.

## Methods

All experiments were conducted in accordance with the UK Animals Scientific Procedures Act (1986) following Home Office Guidelines. Mice were bred from transgenic lines that expressed the genetically encoded calcium indicator GCaMP in excitatory neurons. One mouse expressed GCaMP6f in glutamatergic neurons (double transgenic Ai95(RCL-GCaMP6f)-D × Slc17a7-IRES2-Cre-D). Five mice expressed GCaMP6s in Camk2a-positive (excitatory) neurons (double transgenic tetO-6GCaMP6s × Camk2a-tTA; Wekselblatt et al., 2016) with GCaMP. In a previous survey, neither line was found to display aberrant activity in the form of interictal spikes (Steinmetz et al. 2017).

### Surgery

Surgical procedures were performed under aseptic conditions and under general anaesthesia. Mice were anesthetized with isoflurane (Merial) at 3–5% for induction, and 0.75–1.5% subsequently. Body temperature was maintained at 37 C using a heating pad. Carprofen (5 mg/kg, Rimadyl, Pfizer) was administered subcutaneously for systemic analgesia, and dexamethasone (0.5 mg/kg, Colvasone, Norbrook) was administered to prevent brain swelling. The scalp was shaved and disinfected, and a local analgesic (Lidocaine, 5% ointment, TEVA UK; or intradermal injection, 6 mg/kg, Hameln Pharmaceuticals Ltd) was applied prior to the incision. The eyes were covered with eye-protective gel (Viscotears, Alcon; or Chloramphenicol, Martindale Ltd). The animal was positioned in a stereotaxic frame (Lidocaine ointment was applied to the ear bars), the skin covering and surrounding the area of interest was removed, and the skull was cleaned of connective tissue. A custom headplate was positioned above the area of interest and attached to the bone with Superbond C and B (Sun Medical). Then, a round craniotomy (3–4 mm diameter) was made over the right posterior cortex with a fine-tipped diamond drill and/or a biopsy punch (Kai Medical). The craniotomy was centered at stereotaxic coordinates −2 mm Posterior to Bregma and 2 mm Lateral. The craniotomy was covered with glass coverslip (a 5-mm diameter outer coverslip glued to a 4-mm inner coverslip). A circular metal headplate of 7 mm radius was attached with dental cement. All recordings were thus made in the right posterior cortex only. Following surgery, mice were placed in a heated container until they were ambulatory. Mice were then given carprieve in water as an analgesic and were given at least three days to recover.

### Habituation

Following recovery, mice were habituated gradually to the apparatus. They were first handled in their home cage, then gradually introduced to longer periods of head fixation. Once they were comfortable on the rig, two-photon imaging and wide-field retinotopy was acquired to ensure adequate imaging quality. If these criteria were passed, mice were water restricted so that water could be used as a reward. Body weight was monitored to ensure mice maintained at least 80% of their initial body weight. A minimum water allowance of 40 mL/kg per day was provided. If a mouse did not receive this daily allowance when performing the tasks, the rest of the fluids were delivered afterwards in the form of water or hydrogel. After at least two days of water restriction to ensure a stable weight and no adverse effects, mice were slowly introduced to elements of the task.

### Apparatus

The mouse was head-fixed and surrounded by three computer screens (Iiyama ProLite E1980SD) at right angles, with the central screen ∼20 cm away. The screens spanned ∼270 deg horizontally and 70-75 deg vertically and refreshed at 60 Hz. Fresnel lenses were mounted in front of the screens to correct for aberrations in luminance and contrast at steeper viewing angles, covered with a scattering window film to prevent specular reflections (Burgess et al. 2017). A nearby speaker played auditory stimuli associated with task events, i.e. onset tones, reward tones, and incorrect noise bursts. A water spout was positioned near the mouth. Water delivery was controlled by a valve muffled in a block of foam, which retained an audible click on reward delivery.

### Training

Head-fixed mice were trained on two variants of a two-alternative forced-choice visual detection task for water reward. Both tasks were performed under the same imaging rig. One task involved virtual navigation by running on a styrofoam ball, and the other involved turning a steering wheel to move a visual grating. Mice were usually trained to asymptotic performance on one task before they were introduced to the other. Some mice started with the T-maze and others started with the steering-wheel task.

Both tasks involved a vertical grating on either the left (overlapping with −30 deg azimuth) or right (overlapping with +30 deg azimuth) side of the visual field, at central elevation. In both tasks, mice had to orient in the same direction to bring the stimulus to the center of their visual field to make a correct response. Orienting in the opposite direction, pushing the stimulus away from the center, was an incorrect response. On each trial, a grating was uniformly randomly chosen among 0%, 6%, 12%, 25% and 50% contrasts. Mice received a reward (2 μl of water) for correct choices and a short auditory noise burst for incorrect choices. Both tasks shared task cues such as the onset tone, reward tone, and an initial “interactive delay” of at least 200 ms when movements of the apparatus did not move the stimulus. A gray screen was presented during the inter-trial interval. To help with shaping, in early training, contrasts were initially restricted to including only high contrast subsets and 100% contrast, and mice received larger rewards (3-4 μl). Some mice received sucrose water in training to make the reward more appealing. A shorter ITI was also employed to prevent disengagement by waiting too long in between trials.

### Steering-wheel task

The steering-wheel task is described in Burgess et al. (2017). In the task, mice sit on a raised platform within a “half-pipe” well. Their forepaws rest on a Lego steering wheel which they are able to freely turn in one dimension (rotating in a clockwise or counter-clockwise direction). Stimulus presentation was delivered using the “Rigbox” package (Bhagat et al., 2020). At the beginning of a trial, a visual grating in a Gaussian window (a Gabor stimulus) appears on the left or right side of the screen at ± 30 deg azimuth. The mouse is able to move the wheel to move the grating along the horizontal direction, either an additional 30 deg to the periphery (± 60 deg azimuth) or 30 degs to the center (0 deg azimuth). The wheel was allowed to move immediately, but for the first 200 or 500 ms the stimulus was immobile regardless of wheel movements (interactive delay). Stimulus size was 9 deg (σ of the Gaussian envelope) in initial experiments and 20 deg in later experiments. The inter-trial interval (ITI) was typically 0.5-3 s. The response window was typically 60 s, which was designed to terminate a trial only if mice were extremely inattentive. Wheel movements were detected offline using the “findWheelMoves3” function (github.com/cortex-lab/wheelAnalysis/blob/mast er/+wheel, Steinmetz et al., 2019).

### T-maze task

The T-maze task is described in Krumin et al. (2017). In the task, mice run on a styrofoam ball (20cm in diameter) that is lightly suspended by pressurized air. Movements of the ball were measured using two optical computer mice to control a virtual reality scene. Mice control the ball by running on it. The rotation around the horizontal left-right axis (pitch) was responsible for forward movement in virtual reality, and the rotation of the ball around the vertical axis (yaw) was responsible for turning in virtual reality. The lateral displacement of the ball (rotation around the horizontal front-back axis, roll) was ignored. At the start of each trial, mice are shown a virtual reality T-maze with a long corridor, and two directions to turn at the end perpendicular to the initial corridor. The visual stimulus (grating) was displayed on the entire left or right wall of the initial corridor. The interactive delay was 200 ms. To make their choice, mice needed to run down the initial corridor and turn left or right down the arms of the T, at which point the trial would end and they would receive a reward for a correct choice or auditory white noise burst for an incorrect choice. Trials were separated by an ITI of at least 1.5-3 s. The virtual scene was controlled using a custom virtual reality engine implemented in Matlab utilizing OpenGL through the Psychophysics Toolbox (Kleiner et al., 2007). The initial corridor was 110 cm long including the juncture of the T, and 20 cm wide, with the two arms of the T spanning 60 cm in width, i.e. an additional 20 cm to the left and right. Noise textures were displayed at 20% contrast on the walls and at 40% on the floor. The grating was superimposed additively on the noise texture.

### Testing protocols

Mice were tested serially in two blocks, with full performance of one task before performance on the other. This was necessary to switch the apparatus across tasks. The gap between tasks was usually no more than a few minutes. To obtain similar numbers of trials in each task, mice were switched to the other task when they reached approximately half of their daily water allowance, typically 100-300 trials depending on performance. The second task was stopped when mice reached their minimum daily water allowance, or stopped performing trials or made many consecutive errors, whichever came first. Mice typically performed 100-300 trials/task/session with a duration of 20-60 min per task. Mice were usually tested on the T-maze first, but occasionally the order was switched. All mice included in the dataset had fully learned and reached asymptotic performance in both tasks. To plot behavioral performance, psychometric curves were fitted using maximum likelihood estimation (Busse et al. 2011).

### Passive conditions

In the same imaging session as the tasks, we occasionally imaged the same neurons in a passive condition on the same apparatus as either task, i.e. the spherical treadmill or steering wheel apparatus. These passive conditions usually immediately followed each task, for 5-60 minutes, but occasionally were included immediately before each task instead. During these passive recordings, the screen was uniform gray.

### Hybrid task

In the hybrid task, the mouse uses the apparatus of the steering-wheel task but views the virtual scene of the T-maze. The mouse viewed the location in the middle of the initial corridor (z = 50 cm), and started the trial looking straight ahead (theta = 0 deg). Turning the steering wheel rotated the view angle (theta). The gain of the steering wheel was matched to the original steering-wheel task.

### Widefield imaging

To identify parietal cortex, we mapped known retinotopic areas by presenting sparse visual noise under widefield imaging. The protocol for widefield imaging followed standard procedure from the literature (e.g. Garrett et al., 2014; Zhuang et al, 2017). The entire 4 mm cranial window was imaged under a widefield macroscope with dual illumination using a sCMOS camera (PCO Edge 5.5). Illumination was generated using an LED (Cairn OptoLED) using alternating frames of violet (405 nm, excitation filter ET405/20x) and blue (470nm, excitation filter ET470/40x) light (at 35 Hz each) to capture calcium-dependent fluorescence and calcium-independent hemodynamic activity respectively. The visual stimulus consisted of black and white squares appearing asynchronously on a gray background. Widefield imaging movies were processed to filter out potential hemodynamic artefacts at the “heartbeat” frequency 7-13 Hz. Then a visual field sign map (Sereno et al. 1994; Garrett et al., 2014) was generated by taking the difference (sine of the angle) between the gradients of the azimuth and elevation maps for every pixel. Sign reversals in the gradient maps correspond to traversals across visual areas, which has been useful in locating visual areas across species (Sereno et al. 1994). The target for parietal cortex was chosen as a region overlapping with area A/RL and adjacent to primary visual cortex (V1) as mapped using wide-field imaging above. This region is defined as a parietal area according to the Allen Mouse Brain Common Coordinate Framework (Wang et al., 2020).

### Two-photon imaging

Two-photon imaging was performed in the target location using a ThorLabs B-Scope with a Nikon 16x 0.8 NA water immersion objective. A Ti:Sapphire (Chameleon Ultra II, Coherent Inc.) laser provided excitation at 920nm, with depth-adjusted power level controlled by an electro-optic modulator, i.e. Pockels cell (M350-80LA, Conoptics Inc.). A custom metal cylinder, cone and black cloth was used to prevent light contamination from the illuminated screens. Acquisition was controlled using ScanImage (Pologruto et al., 2003), and frames were acquired continuously at 30 Hz over an imaging window of 500×500 μm, at a resolution of 512×512 pixels. Multi-plane imaging was performed using a piezo motor over two planes in layer 2/3, starting at 90-130 μm below the surface, separated by 60-70 μm, spanning a total of 180-210 μm. The effective imaging rate was 10 Hz per plane (the flyback plane was discarded).

At the beginning of each acquisition, the mean image over several frames of the previous task’s recording plane was used as a reference plane, and the “live” movie of the current imaging plane was manually aligned in z, x, and y, to match until the difference was indistinguishable to the eye. Following acquisition, the raw movies were then examined by eye, with particular attention before and after the switch in task, to ensure that the same population of cells was visible. Imaging sessions were dropped if a large proportion of neurons were no longer visible by the end of each task or across both tasks. This realignment procedure was most important when switching between the tasks, and also helped the alignment between sessions on different recording days. This procedure was not needed for the days when steering wheel and hybrid task acquisitions were performed, as the mouse stayed with the same head fixation throughout the recording session.

The movies comprising all conditions within a given imaging session (the two tasks plus other conditions) were concatenated before processing in suite2p. Raw movies were processed in Suite2p (Pachitariu et al., bioRxiv) for motion correction (registration), cell detection, signal extraction, neuropil correction and spike deconvolution. Neuropil was estimated as a radius of size 5x the number of pixels defined for the cell and subtracted from cell activity using a multiplicative coefficient estimated per cell, usually ∼0.6-0.8. Deconvolution was performed using the OASIS algorithm (Friedrich et al. 2017) wrapper within Suite2p. Regions of interest (ROIs) detected by Suite2p were manually curated using the Suite2p Graphical User Interface. ROIs were classified as cells according to spatial and temporal criteria, i.e. that the ROI reasonably resembled a disc-like soma at the size expected at the imaging zoom used and that the inspected activity trace had good signal-to-noise. Manual curation was performed blind to the time at which the task transition occurred.

To judge consistency of the results, we returned to the same cells across days, using RegisterS2p to align recorded ROIs across days and identify matches (Pachitariu et al. 2018). We only analyzed neighboring pairs of days (separated by one or two days) as this ensured that recorded cells were most similar, with respect to morphology, cell death and changing neural representations across longer timescales if any. Pairs of sessions were upheld to the same strict criteria for inclusion as described above, so n = 4 pairs of days remained for analysis. The cells analyzed were the union of cells present in each pair, and we analyzed pairs of task conditions, either the same task across days, or different tasks across days.

### Quantifying neural activity

All analyses were carried out on a session-by-session basis. Summary statistics were then taken across sessions. To summarize average neural activity in the whole session we used “isolation distance” (Stringer & Pachitariu, 2019). This measure captures deviations of a cell’s activity relative to its neuropil surround. Specifically, for each cell and its respective neuropil surround (both estimated in Suite2p) the matrix of pixels × time is concatenated over pixels, and the mean (over both cell and neuropil) is subtracted over time. Then singular value decomposition is used to reduce the dimensionality to the first principal component, resulting in a one-dimensional summary per pixel. The “distance” between the distributions of the pixels corresponding to the cell and neuropil is then computed. Specifically, we used the Bhattarcharya distance, which accounts for the variance of each distribution, since the neuropil distribution tends to have less variance (over the range spanned by the cell).

In non-soma-localized GCaMP indicators, GCaMP is present not just in the cell bodies but also dendrites and passing axons. Out-of-focus fluorescence from this “neuropil” can erroneously contribute to the signal averaged within the pixels that define a cell. A standard procedure is to correct for this neuropil by subtracting a scalar multiple of the average activity in a radius around each cell (e.g. Chen et al., 2013; Dipoppa et al., 2018). Here the “neuropil coefficient” was estimated per cell but is usually <1, around ∼0.6-0.8. Meanwhile, standard cell extraction procedures for two-photon data involve estimating pixels which are correlated within themselves but not with respect to the surrounding pixels in the background, i.e. the neuropil. Given these are well-established assumptions in the literature about what constitutes a cell and what constitutes extraneous noise to be subtracted out, it is reasonable to assume the neuropil can be treated as an estimate of baseline “noise”. Isolation distance uses this assumption to compute single-neuron “activity” as the difference between activity within the ROI and activity in the neuropil surround — in effect a measure of signal-to-noise ratio. There is some precedent for this approach as applied to calcium imaging (Chen et al., 2015).

Isolation distance produced results qualitatively similar to common measures (mean, standard deviation, skewness, coefficient of variation) but importantly was most robust to baseline noise. This last requirement was especially important as a high noise floor is observed in the strain of GCaMP6s transgenic mice that contributed to the majority (5/6 mice) of our dataset (as observed in Huang et al. 2019).

### Running and stationarity

To determine if task selectivity could be explained by running modulation, we used the “passive ball” condition, as running therein provided a task-agnostic condition to assess a neuron’s modulation by running. Running modulation was computed as the correlation between each cell’s activity and the mouse’s running speed. The deconvolved, neuropil-corrected calcium trace was used to account for movement artefacts which can occur due to fast z-drift (Stringer, 2018). The running speed was taken as forward movement on the ball, and was downsampled to match the imaging frame rate. Both the running speed and neural activity were smoothed by convolving the traces with a 1-s s.d. Gaussian filter. A permutation test was used to assess significance by circularly shifting running speed relative to the neural activity 1,000 times by a random number of frames.

We further tested whether isolation distance during the steering-wheel task reflected a qualitatively distinct state of stationarity, that was independent from the running state in the T-maze. Blank screen recordings were often fairly short (usually 10-20 minutes), and mice often ran during these, so there were not enough time points when the mouse was stationary to use this condition. Instead, we used stationary periods in the T-maze. We re-computed isolation distance in the T-maze only including the frames when the mouse had a running speed <1.2cm/s and was stationary for at least 3s — the latter to avoid contamination by preceding running periods due to slow decay of GCaMP, and also to ensure the stationarity was not a brief pause during a change in angular velocity. If isolation distance is a mere function of running vs stationary states, we would expect isolation distance to be highly positively correlated, as these conditions now belong to the same motor state (stationary).

### Choice selectivity

To determine choice selectivity we used the mean deconvolved calcium activity over the whole trial, from stimulus presentation and including the motor execution of the choice. In some sessions, the same stimulus condition was repeated if the mouse did not respond correctly, to encourage engagement. These repeated trials were excluded from analyses, as mice could know with certainty the correct choice even prior to the trial, and thus may engage in a different strategy for choices that is not guided by sensory evidence. Trials in which the wheel moved <125 ms after stimulus onset were discarded as such movements are unlikely to be a response to the stimulus. Sessions were only included if at least 10 trials of each comparison (e.g. 10 left-side choices and 10 right-side choices) remained after excluding these invalid trials defined above. For analyses comparing choice selectivity across tasks, there needed to be 10 trials for each choice and each task for a session to be included.

In well-performing mice, stimulus and choice are highly correlated; to disentangle these factors and focus on choice alone, we used a measure called “combined conditions choice probability” (ccCP) introduced elsewhere (Steinmetz et al. 2019). Like original measures of choice selectivity (Britten et al., 1992), this measure only compares neural activity for different choices within the same stimulus condition. However, ccCP is able to use all trials from the full stimulus set (nine contrast conditions), and therefore provides a better estimate of choice selectivity when there are few trials for individual contrast conditions. We normalized the ccCP to lie between −1 and 1, where negative values mean higher activity during left choices. To assess the significance of computed choice selectivity values we used a permutation test. For every neuron, trial labels were shuffled 1,000 times, and choice selectivity was recomputed for each new batch of pseudo “left”-labelled and “right”-labelled trials. As the same number of trials remain in each comparison, the permutation test accounts for imbalanced samples of each condition. To compare choice selectivity across tasks, we only used the subset of neurons active in both tasks, chosen by a threshold of an isolation distance of > 0.3. Sessions with 10 or fewer neurons active in both tasks were excluded.

## Supplementary figures

**Supplementary Figure S1.**
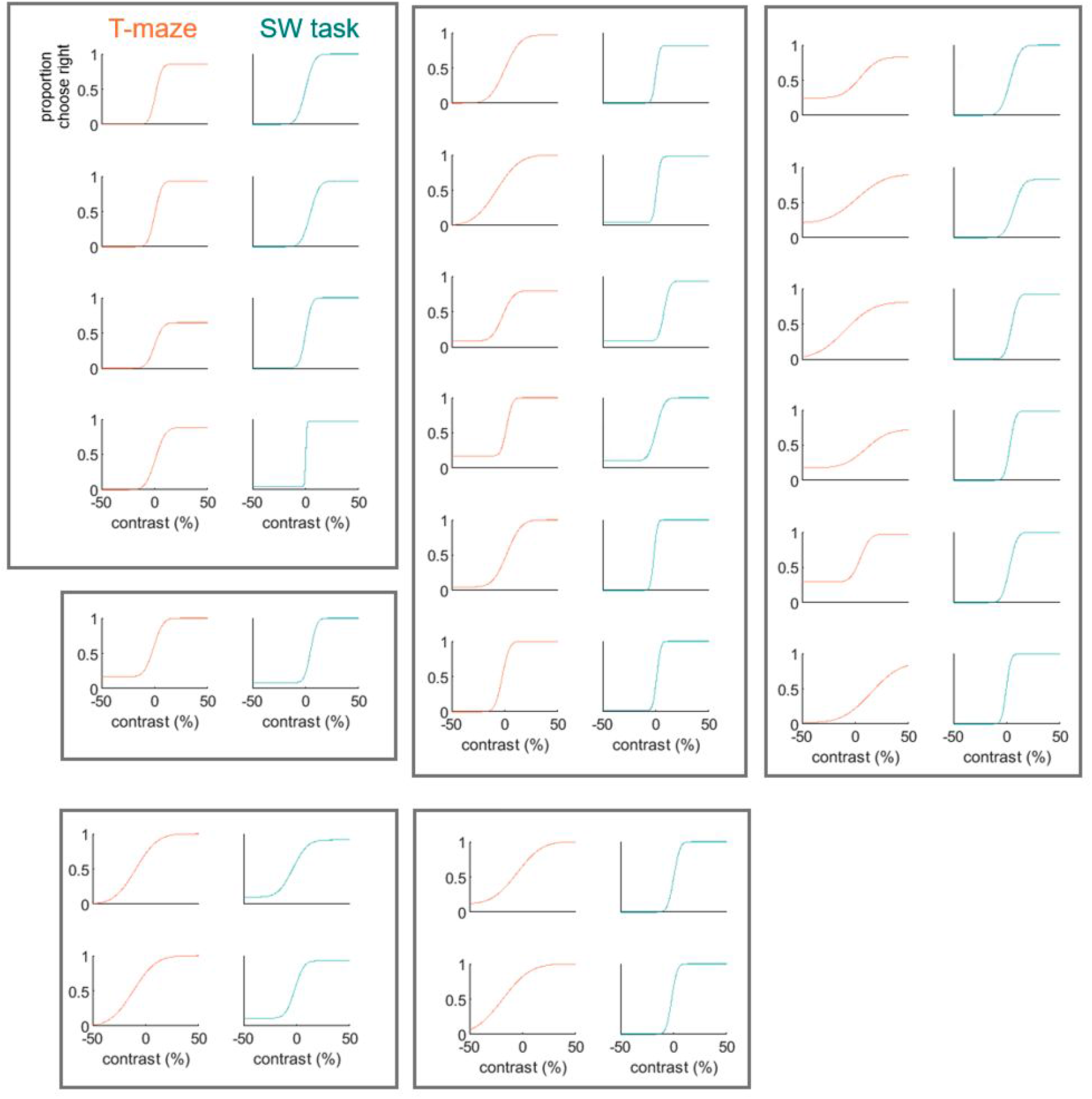
Psychometric fits for each task in all sessions in six mice. Each box shows the collection of sessions belonging to a different mouse. Each row within a box is a single session where both tasks were performed consecutively in the same day. Tasks were not necessarily presented in the order shown. All included sessions are after mice had fully learned both tasks.

**Supplementary Figure S2.**
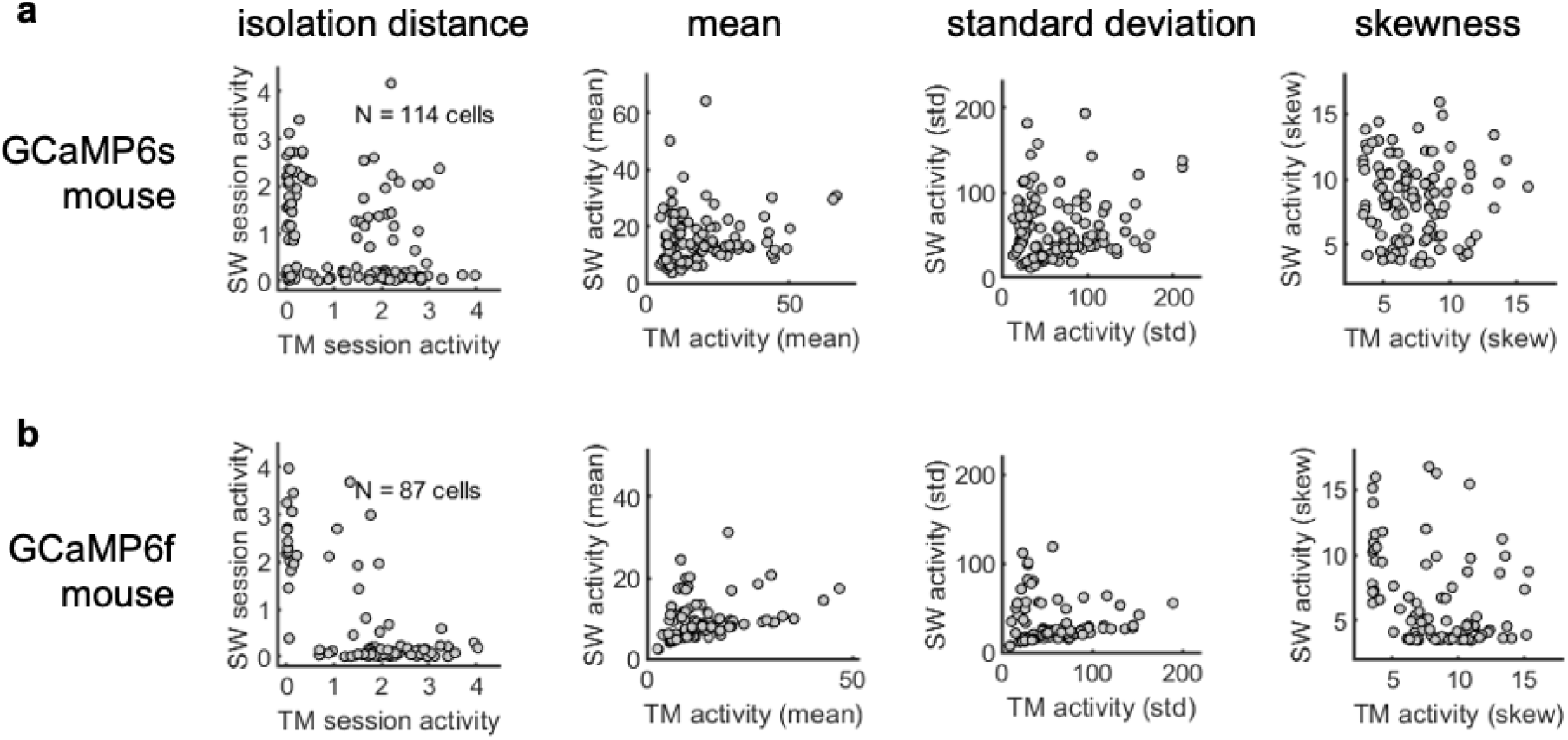
Comparison of activity measures. Example sessions comparing different measures for calculating activity: isolation distance, mean, standard deviation and skewness. (a) Example session from a GCaMP6s mouse. (b) Example session from a GCaMP6f mouse.

**Supplementary Figure S3.**
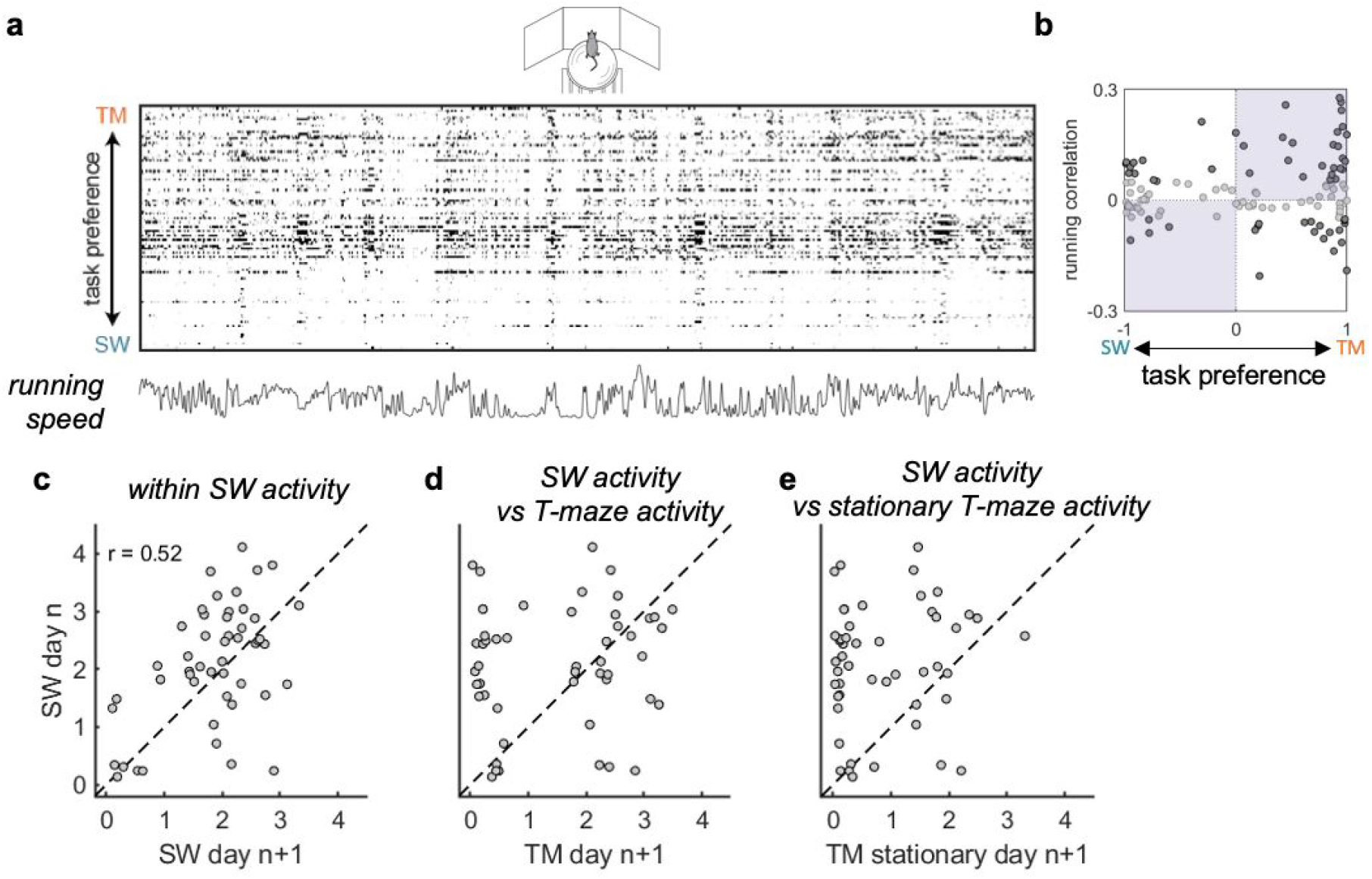
Running modulation does not account for task specificity. (a) Firing rate of neurons from an example session during the passive treadmill condition, sorted by task selectivity. Below: simultaneously-recorded running speed of the mouse. (b) Running modulation as a function of T-maze vs steering-wheel task preference (positive values mean T-maze preference). Filled circles show significant running-modulated neurons (permutation test). The purple overlay shows hypothetical distributions of cells, if T-maze neurons were solely those modulated by running, and steering-wheel task neurons were solely those suppressed by running. In this example session, the Spearman rank correlation was not significant, using all neurons, r = 0.04, p = 0.67, or just significantly running-modulated neurons (filled circles), r = 0.06, p = 0.65. (c) Comparison of steering-wheel task activity across days in two example sessions, r = 0.52, p < 1e-3. (d) Same example sessions, comparing across tasks; activity is not significantly correlated, r = 0.13, p = 0.39. (e) Comparison of steering-wheel activity to only stationary periods in the T-maze; activity is not significantly correlated, r = 0.12, p = 0.41.

## Notes

### Competing Interest Statement

The authors have declared no competing interest.

